# Spatial neglect in the digital age – Influence of presentation format on patients’ test behavior

**DOI:** 10.1101/2021.06.11.447882

**Authors:** Hannah Rosenzopf, Christoph Sperber, Franz Wortha, Daniel Wiesen, Annika Muth, Elise Klein, Korbinian Möller, Hans-Otto Karnath

**Affiliations:** Centre of Neurology, Division of Neuropsychology, Hertie-Institute for Clinical Brain Research, University of Tübingen, Tübingen, Germany; Department of Psychology, University of Greifswald, Greifswald, Germany; University of Paris, LaPsyDÉ, CNRS, Sorbonne Paris Cité, Paris, France; Leibniz Institut für Wissensmedien, Tübingen, Germany; Centre for Mathematical Cognition, School of Science, Loughborough University, United Kingdom; Centre for Individual Development and Adaptive Education of Children at Risk (IDeA), Frankfurt, Germany; Department of Psychology, University of South Carolina, Columbia, SC

**Keywords:** Diagnostics, Test digitalization, Center of Cancellation, Right hemisphere, Stroke, Human

## Abstract

Computerization of diagnostic neglect tests can deepen our knowledge of neglect specific abnormalities, by effortlessly providing additional behavioral markers that are hardly extractable from existing paper-and-pencil versions. However, so far it is not known whether the digitization and/or a change in size format impact neglect patients’ search behavior and test scores and thus require adjustments of cut-off criteria. We compared the Center of cancellation (CoC) measure of right hemisphere stroke patients with spatial neglect in two cancellation tasks across different modalities (paper-and-pencil vs. digital) and display sizes (small, medium, large). We found that the CoC measure did neither vary considerably between paper-and-pencil versus digital versions, nor between different digital size formats. The CoC derived from cancellation tasks thus seems robust to test digitization. A further aim of the present study was to evaluate three additional parameters of search behavior which became available through digitization. We observed slower search behavior, increased distance between two consecutively identified items, and signs of a more strategic search for neglect patients than control patients without neglect. Machine learning classifications indicated that – beyond the CoC measure – the latter three variables can help to differentiate stroke patients with spatial neglect from thosewithout.

## 1. Introduction

Spatial neglect is a common and debilitating consequence of brain damage mainly to the right hemisphere (Becker & Karnath, 2007). Core symptoms of neglect − as defined by Corbetta and Shulman (2011) and Karnath and Rorden (2012) − include an egocentric bias in gaze direction and exploration towards the ipsilesional side. One type of test to detect and quantify these core symptoms are cancellation tasks (Weintraub & Mesulam, 1985; Gauthier, Dehaut & Joanette, 1989; Ferber & Karnath, 2001). Such tasks are commonly presented on sheets of paper placed in front of the patient, who is required to find and manually mark all targets among distractors. Patients with spatial neglect often miss targets on the contralesional side. The presence and severity of spatial neglect can be measured by computing the Center of Cancellation (CoC, Rorden & Karnath, 2010) which assesses the average position of correctly marked targets with respect to the patient’s egocenter.

While paper-and-pencil based cancellation tasks can be a time-efficient yet reliable diagnostic alternative to more extensive test batteries (Fullerton, Stout, & McSherry, 1986; Ferber & Karnath, 2001), they provide only part of the information they could if they were computer-based (Schendel & Robertson, 2002; Bonato & Deouell, 2013, Dalmaijer, Van der Stigchel, Nijboer, Cornelissen, & Husain, 2015). Among other aspects, digitization can provide additional variables such as, e.g., response time, revisits (of already marked items), and information concerning the search path applied (Donelly et al., 1999; Dalmaijer et al., 2015). However, due to the lack of comparison with the traditional, validated paper-and-pencil version, it cannot yet be ruled out that variations in modality may lead to results that differ from those of traditional paper-and-pencil versions.

Furthermore, in clinical practice, traditional A4 paper-and-pencil tests will likely be implemented as scaled-down versions to match commonly used tablet formats. However, the effect of using devices of different sizes on the validity of the tests has not yet been sufficiently studied in the context of cancellation tasks. Previous observations have suggested that the size or − more precisely − the length of the stimulus material may have some influence on spatial attentional processing (Bowers & Heilman, 1980; McCourt & Jewell, 1999; Anderson, 1997; McCourt & Jewell, 1999). On the other hand, studies in neurological patients have suggested that a change in frame size does not necessarily affect neglect-specific impairments. Body-centered (egocentric) and object-centred (allocentric) neglect appeared to dynamically adapt to different frame sizes (Karnath & Niemeier, 2002; Baylis, Baylis, & Gore, 2004; Karnath, Mandler & Clavagnier, 2011; Li, Karnath & Rorden, 2015).

In the present study, we compared right hemisphere stroke patients’ performance in cancellation tasks across different modalities (paper-and-pencil vs. digital) and display sizes (small, medium, large) to investigate whether digitization of traditional cancellation tasks to various screen sizes affects their validity. As new variables become available through digitization, a further objective was to evaluate these variables with respect to their contribution to diagnostic decisions. This is important, because in clinical practice it often happens that patients cannot complete a cancellation task (e.g. because they are too exhausted or because testing must be interrupted due to other clinical necessities). While measuring the CoC requires running the test to completion, the behavioral aspects measured by these new variables might become extractable already early on and thus aid diagnosis (if a test cannot be completed), given that these new parameters proved to detect neglect specific behavior. The latter shall be investigated in the following. Further, methodological investigations have shown that effects revealed by statistical analyses often have limited informative value in (applied) diagnostic contexts, even when effect sizes are very large (Dwyer, Falkai & Koutsouleris, 2018). Machine learning analyses are a particularly promising approach to address these issues and reveal the diagnostic value of new variables and measures. Accordingly, we decided to test the potential diagnostic value of process parameters obtained from digital cancellation tests using such methodological approaches.

## 2. Materials and Methods

### 2.1 Subjects

Twenty continuously admitted right hemisphere stroke patients (N=12 suffering from spatial neglect; N=8 without spatial neglect) were recruited at the Centre of Neurology at Tuebingen University. Structural imaging was acquired by computed tomography (CT) as part of the clinical routine procedure carried out for all stroke patients at admission except for one patient who received magnetic resonance imaging instead (MRI). Patients with diffuse or bilateral brain lesions, patients with tumors, and patients in whom scans revealed no obvious lesions were not included. Clinical and demographic variables of the two patient groups are summarized in Table 1; Figure 1 illustrates a simple overlap plot of their brain lesions. The study was performed in accordance with the revised Declaration of Helsinki, the local ethics committee approved the study and all patients provided their written consent to participate.

**Table 1.** Demographic and clinical data of the 20 right hemispheric patients with and without spatial neglect included in the study. Mean, standard deviation.

|  | Neglect | No neglect |
| --- | --- | --- |
| Age in years | 71.0, 8.3 | 56.9, 15.9 |
| Sex (M/F) | 8/4 | 7/1 |
| Etiology (Ischemia/Hemorrhage) | 12/0 | 8/0 |
| Visual Field Defects (% present) | 25 | 12.5 |
| Hemiparesis (% present) | 100 | 78 |
| Letters cancellation test (CoC) | 0.330, 0.419 | 0.0069, 0.0336 |
| Bells test (CoC) | 0.415, 0.252 | - 0.0079, 0.0461 |
| Copying task (N omitted) | 2.56 | 0.38 |

**Figure 1:**
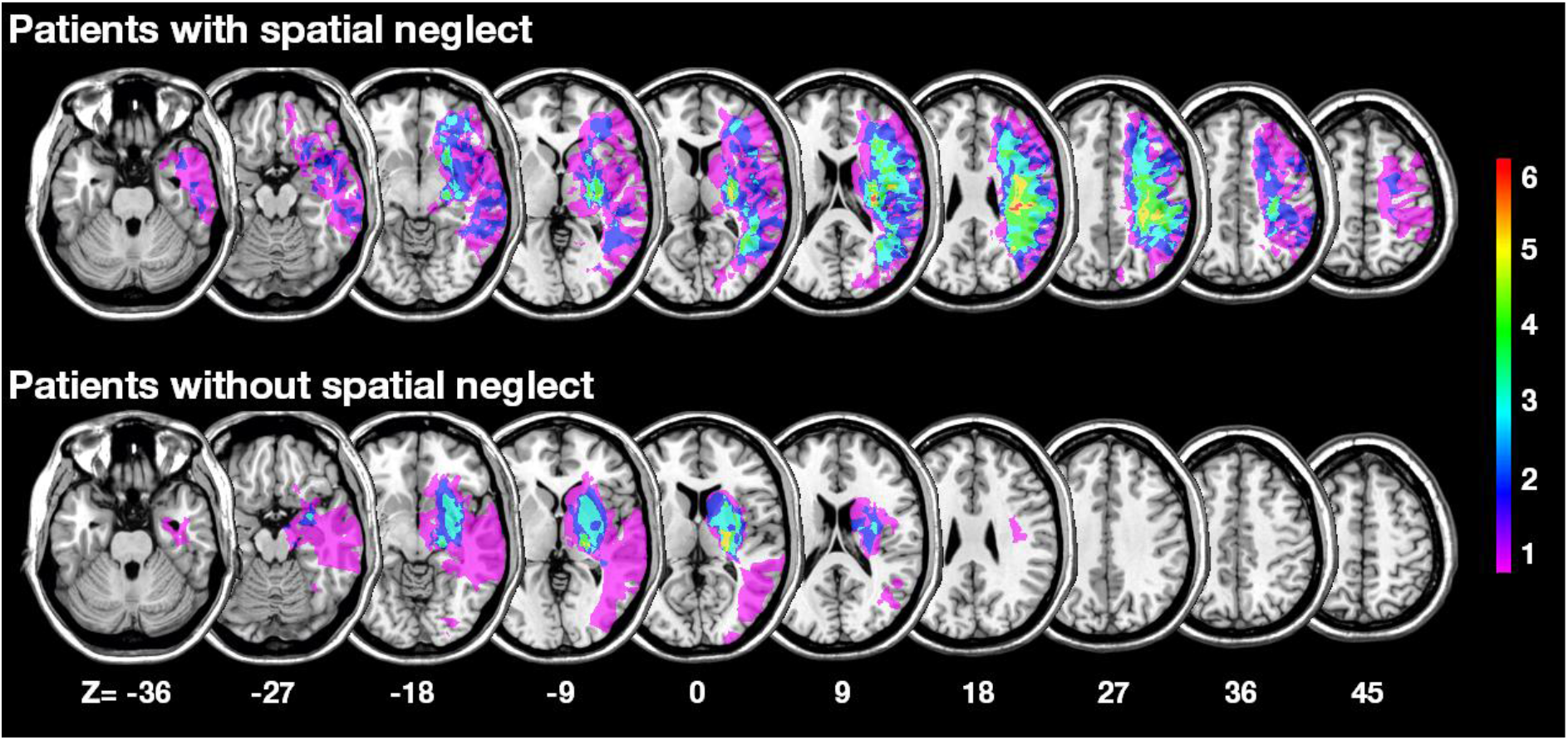
Lesion overlays. Overlay of the normalized lesions of right hemisphere patient groups with and without spatial neglect. Lesion boundaries were semi-automatically using the Clusterize algorithm on the SPM Clusterize toolbox (cf. de Haan et al., 2015) on SPM 12 (www.fil.ion.ucl.ac.uk/spm). Normalization of CT or MR scans to MNI space with 1×1×1 mm resolution was performed by using the Clinical Toolbox (Rorden et al., 2012) under SPM12, and by registering lesions to its age-specific MR or CT templates oriented in MNI space (Rorden et al., 2012).

All patients were clinically examined with a bedside neglect screening after being first admitted to the Centre of Neurology. Nineteen patients were tested in the acute phase (on average 6.4 days, *SD* = 4.5 post-stroke); one patient from the neglect group was tested in the chronic phase (32 months post-stroke). The screening included two cancellation tasks (Bells test [Gauthier et al., 1989]; Letter cancellation [Weintraub & Mesulam, 1985]), and a copying task (Karnath et al., 2002). These tasks were presented on a DIN A4 sized 297 by 210 mm paper each. For the Letter and Bells Cancellation tasks, we calculated the Center of Cancellation (CoC) using the procedure and cut-off scores for diagnosing spatial neglect by Rorden and Karnath (2010). The CoC is a sensitive measure capturing both the number of omissions, as well their location. In the copying task, patients were asked to copy a complex multi-object scene consisting of four figures (a fence, a car, a house, and a tree), with two of them located in each half of the horizontally oriented sheet of paper. Omission of at least one of the contralateral features of each figure was scored as 1, and omission of each whole figure was scored as 2. One additional point was given when contralateral located figures were drawn on the ipsilesional side of the paper sheet. The maximum score was 8. A score higher than 1 (i.e. > 12.5% omissions) was taken to indicate neglect. The maximum duration of each test was not fixed but depended on the patient being satisfied with his/her performance and confirming this twice. Patients were diagnosed with spatial neglect if they scored above the cut off in at least 2 out of 3 tests (cf. Tab. 1).

### 2.2 Experimental apparatus and procedure

The experiment included the same cancellation tasks as in the clinical assessment (see above), i.e. the Bells test and the Cancellation task, presented on A4 sheets of paper. Beyond, the experiment comprised computerized touch screen versions of the Bells test and the Cancellation task. Computerized testing was performed on a capacitive 27-inch multi-touch display (3M – M2767PW), connected to a laptop (HP ProBook 4740s with Windows 7 Professional). The touchscreen versions of the two tasks were custom created using MATLAB 2016b and Psychtoolbox (http://dx.doi.org/10.17632/6dzxs69j7d.1). Computerized cancellation tests were high-resolution versions of the original test images used for the paper-and-pencil version, but were displayed in three different tablet sizes (TS): 260.28 mm × 173.52 mm (‘TS small’; a tablet format as e.g. in Microsoft surface, HP Elite, Dell Latitude 5290), 297 mm × 210 mm (‘TS medium’; equivalent to A4 paper sheet size), and 597.6 mm × 336.2 mm (‘TS large’; full-screen size of the 27-inch touch screen). To keep paper-and-pencil and touchscreen conditions as comparable as possible, the touchscreen always lay flat on the table and a touchscreen compatible pen (Adonit Dash 2) was used to mark the targets on it. Patients’ marks were visualized in real-time, providing patients with visual feedback comparable to that provided by conventional pens on a regular sheet of paper. Due to their health issues, four patients were unable to complete all trials, which led to 9 missing data sets in different test conditions and thus had to be excluded from parts of the analyses.

In the experiment, half of the participants started with the paper-and-pencil version of the two cancellation tasks, the other half with the touchpad version. The order of the two paper-and-pencil versions was alternated, the order of the 6 different touchpad versions was randomized. Participants were instructed to find all the bells/letters ‘A’ that were spread around distractors and to tell the experimenter once they were done. Before starting the next trial, all patients were asked to confirm that they were indeed done with this trial, i.e. could not find any other target stimuli.

### 2.3 Data Analysis

For comparing right hemisphere stroke patients’ CoC performance in cancellation tasks across different modalities (paper-and-pencil vs. digital) and display sizes (small, medium, large), we used Wilcoxon and Friedman tests respectively. The new, additional parameters of patients' search behavior that we extracted from the digital test version were (a) the number of targets found relative to time, (b) the mean distance between every two targets found in direct succession to each other, and (c) an indicator for whether or not participants applied a strategy in their search. For (a) targets per time, we extracted a participant’s total number of correctly identified items and divided it by the time measured between starting the test and marking the last item. Next, (b) we averaged the Euclidean distance between every two targets found in direct succession to each other. To investigate (c) whether or not a search strategy was applied when exploring the screen, the distance between each correctly identified target and the consecutively identified target was calculated again. However, this time it was separated into horizontal (x-axis) and vertical (y-axis) distance. Both distances were standardized to the screen size, resulting in a unitless measure, and averaged across all found targets. A strategic search, as we define it here, should result in low values in either the mean x-axis distance or the mean y-axis distance. Low y values indicate a row-wise left-to-right (reading-like; Fig. 2A) or alternating left-to-right and right-to-left (Fig. 2B) search pattern; low x values indicate a column-wise top-to-bottom (Fig. 2C) or alternating top-to-bottom and bottom-to-top (Fig. 2D) search pattern. The measure is independent of direction and applies also if tests were started from the right or the bottom. To investigate potential differences between (i) the digital screen sizes and (ii) right hemispheric patients with spatial neglect in comparison to patients without neglect, we applied a 2×3 analyses of variance (ANOVA) for each of the three parameters above (i.e., time per target, mean distance, and search strategy) with the between-subjects factor group (neglect vs. no neglect) and the within-subjects factor screen format (TS small vs. TS medium vs. TS large).

**Figure 2:**
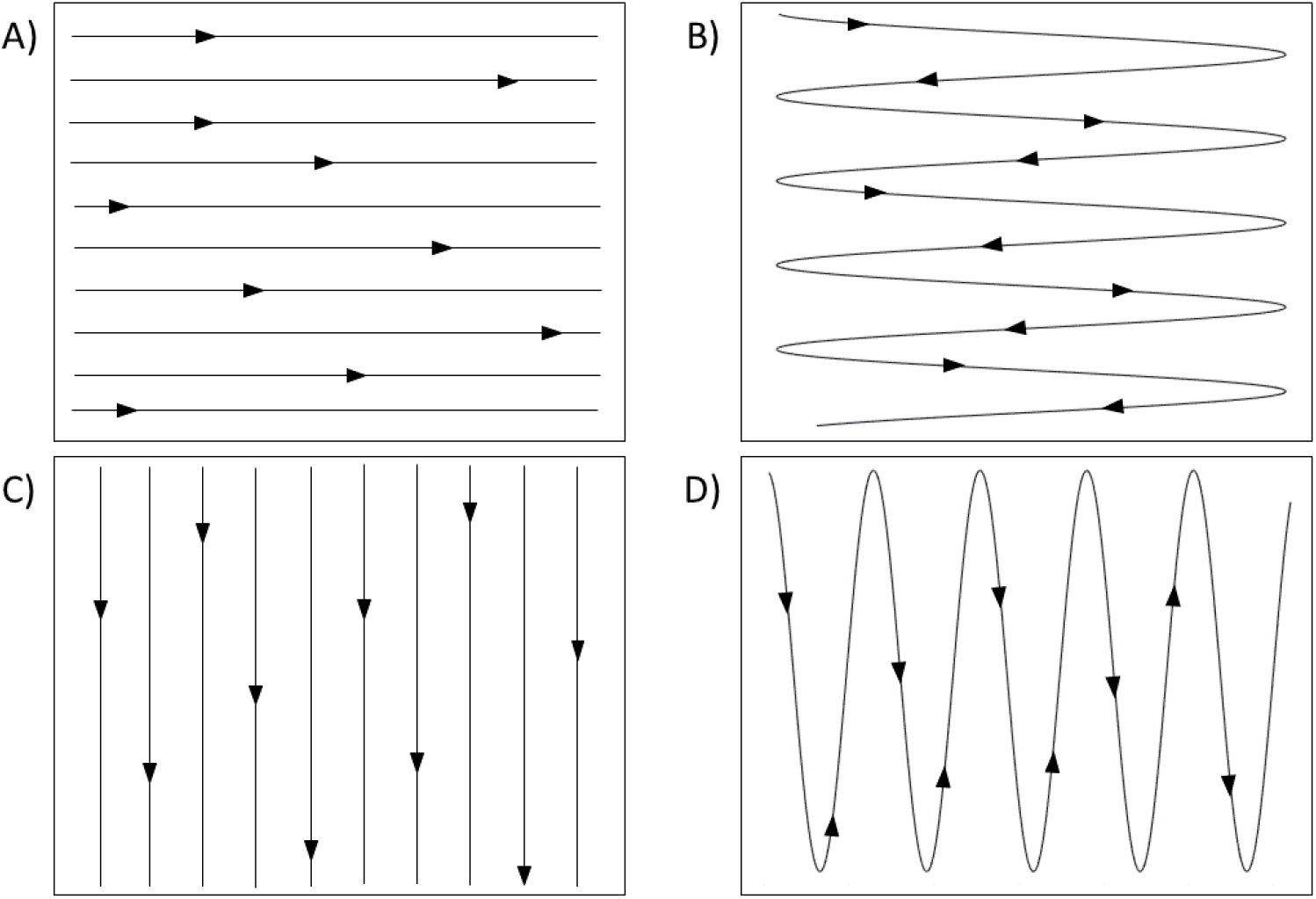
Measured search strategies. Search strategies covered by variable “search strategy” in the present study: (A) left-to-right (reading-like) strategy; (B) alternating left-to-right and right-to-left search pattern; (C) column-wise top-to-bottom strategy; (D) alternating top-to-bottom and bottom-to-top search pattern.

To finally analyze if the three parameters above can be used to reliably predict participants’ neglect diagnosis (dichotomized: spatial neglect vs. no neglect), we used Support Vector Machines (SVM). Given that SVM require complete data sets, we first used Multiple Imputation by Chained Equations (MICE; White, Royston, & Wood, 2011) to impute missing data for this analysis step only. It entailed that missing values in a given column were estimated using a Bayesian Ridge Regressor, predicting values of the current column from all other columns. MICE was carried out column-wise from the column with the least number of missing values to the column with the most missing values. The potential impact of the imputation was tested by re-reruning all analyses with a dataset where missing values were omitted. In the following sections only results for the imputed dataset will be reported, because the pattern of results remained identical with and without imputation. Due to the sample size of the present study, we decided to use a dataset containing each screen size (TS small, TS medium, TS large) for each participant. To account for the dependence of data points in this approach (i.e., having three measures for each participant), we tested our models through Leave-One-Subject-Out Cross-Validation. In this procedure, the machine learning model is trained on data for all participants but one and tested on the participant that was left out for training. This process is then repeated until each participant was predicted once and prediction outcomes (i.e., balanced accuracy due to the unequal group sizes; Brodersen, Ong, Stephan & Buhmann, 2010) are averaged across predictions for all participants. Hyperparameter (i.e., the kernel: linear or radial basis function; cost parameter: ranging from 0.01 to 10) were optimized through a grid search in a nested Leave-One-Subject-Out Cross-Validation (within the training dataset). This procedure was carried out separately for each task (Bells test and Letter cancellation) and balanced accuracy scores were obtained across all screen sizes for both tasks. Lastly, the percent of correctly classified neglect and right-hemispheric control patients were accumulated for each screen size and test. To test if the classification accuracy varied by screen size, chi square tests of independence were used comparing the distribution of correctly classified neglect and right-hemispheric control patients across screen sizes for each task (Bells test and letter cancellation). All machine learning analyses were conducted in Python using the scikit learn module (Pedregosa et al., 2011).

## 3. Results

### 3.1 Comparison between paper-and-pencil and digital formats

To investigate whether digital versus paper-and-pencil test modality has an impact on patients’ performance in cancellation tasks, we compared patients’ mean CoC scores in the A4 paper-and-pencil version to those in the same format in the digital A4 touch screen version (TS medium). Data are illustrated in Figure 3. Wilcoxon tests indicated no significant median CoC differences between the digital and the paper-and-pencil versions, neither in the Letter cancellation test (*Z*= 1.784, *p* = 0.072) nor in the Bells test (*Z*=0.533, *p* = 0.594). In clinical practice, the traditional A4 paper-and-pencil tests will most likely be implemented as a downscaled version to match the currently used tablet format. Thus, we also investigated (cf. Fig. 3) whether differences in performance arise between the established A4 paper-and-pencil version and the digital downsized tablet format (TS small). Again, we did not find significant differences for neither the Letter cancellation test (*Z* = 1.784, *p* = 0.074) nor the Bells test (*Z* = − 0.356, *p* = 0.722).

**Figure 3.**
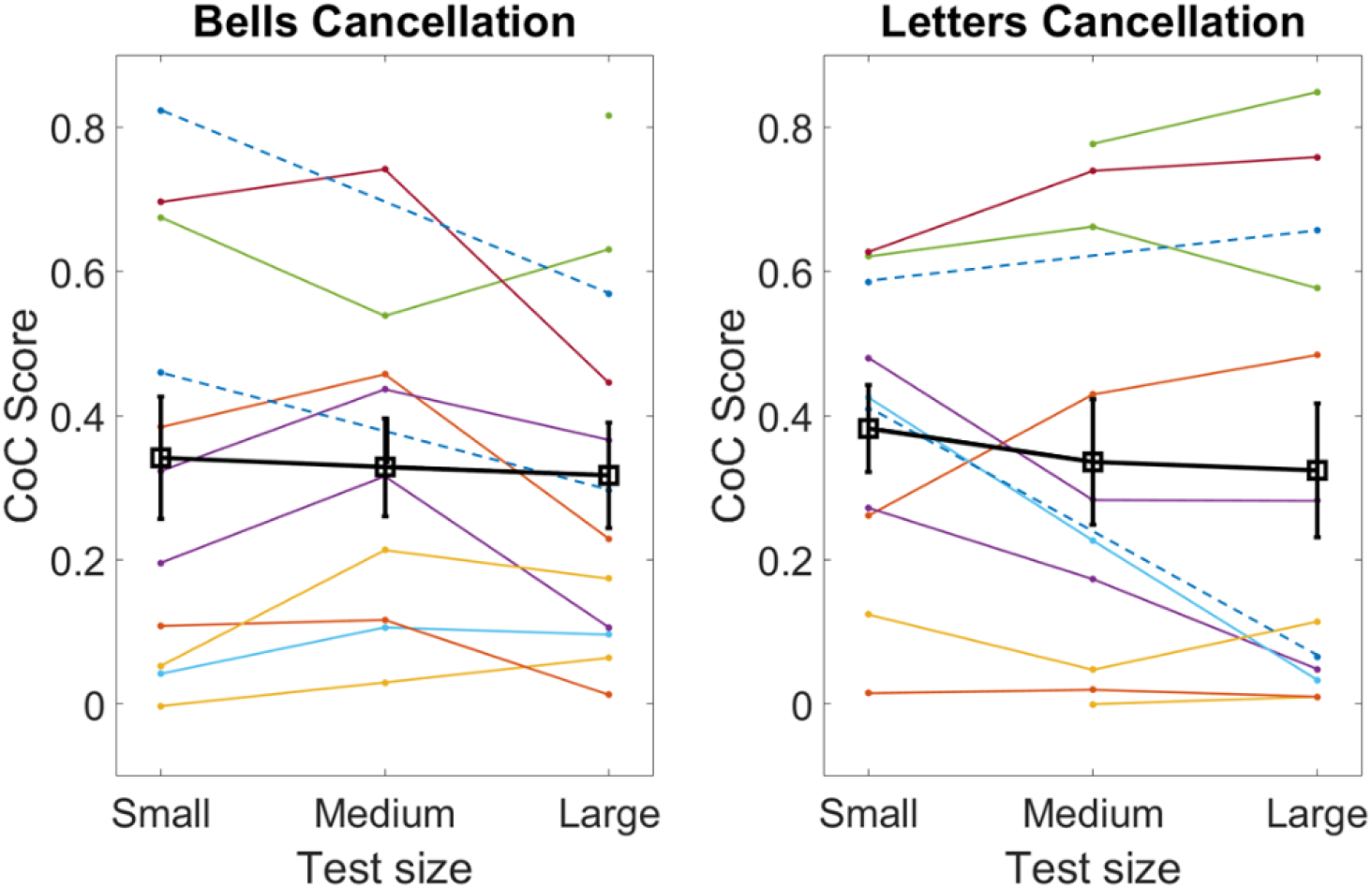
CoC scores over different digital test sizes. Neglect patients’ performance for the Bells test and the Letter cancellation test in the traditional A4 paper-and-pencil version compared to the digital format with an equivalent touchscreen size (TS medium) as well as to the digital format with the smaller size that corresponds to current tablet format (TS small). The bold lines represent mean values (including error bars) averaged over all patients.

### 3.2 Comparison between different sizes of the digital format

#### Center of Cancellation

To investigate whether size variation between the digital versions affects cancellation performances, we used the CoC as dependent variable and test size (TS small vs. TS medium vs. TS large) as independent variable. Data are illustrated in Figure 4. Friedman tests revealed neither significant results for the Letter cancellation test (χ^2^*F*(2) = 1.750, *p* = 0.417) nor for the Bells test (χ^2^*F*(2) = 6.00, *p* = 0.050). Since the latter result was right at the border of significance, we applied post-hoc Wilcoxon comparisons to rule out significant differences. Indeed all three were found to be non-significant.

**Figure 4:**
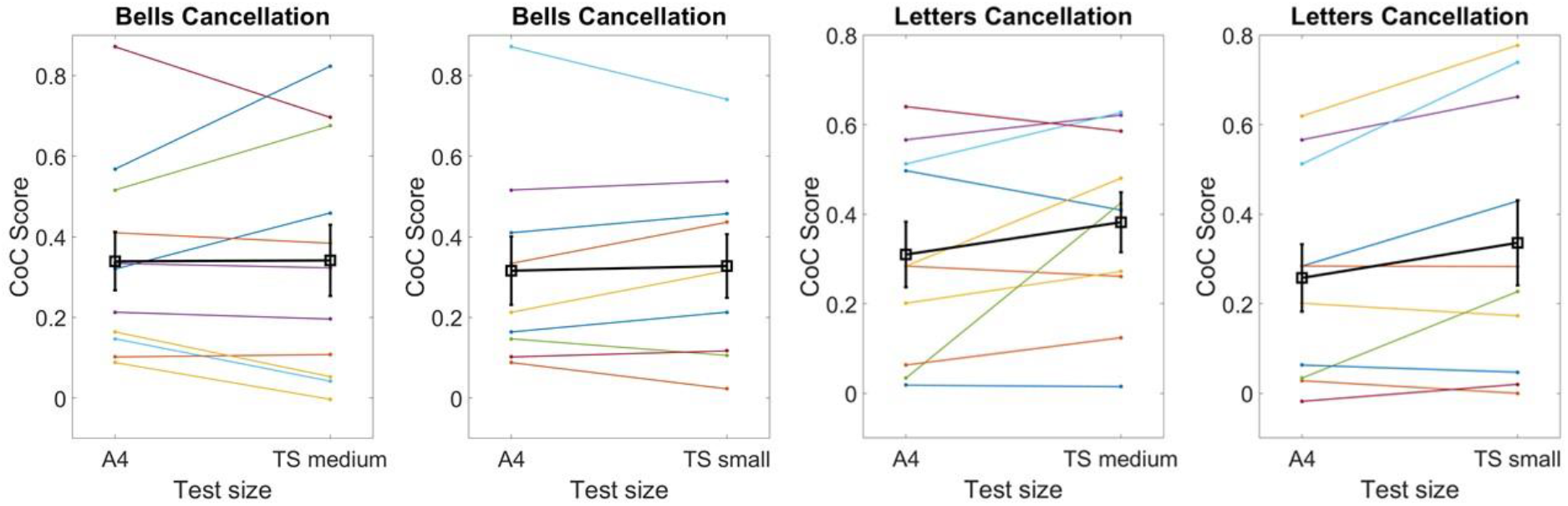
CoC in paper and pencil vs. digital versions. Neglect patients’ performances in the three different touch screen formats of the digitalized Bells test and Letters cancellation test. Test scores of each patient were connected with a line (patients who failed to complete the medium test size version were indicated by a broken line). The bold lines represent mean values (including standard error) averaged over all patients.

#### Additional parameters of search behavior

Beyond the CoC, further variables became available through digitization of the two cancellation tests: targets per time, mean distance, and search strategy.

##### Targets per time

Data are illustrated in Figure 5. Analysis of the Bells test revealed a significant main effect of group (*F*(1,15) = 6.719, *p* = 0.02), indicating that right hemispheric control patients found significantly more targets per time (*M* = 0.196, *SD* = 0.057) than neglect patients (*M* = 0.124, *SD* = 0.066). The main effect of screen size, on the other hand, was not significant (*F*(2,30) = 0.235, *p* = 0.792), indicating that a comparable number of targets was found per time in all three screen sizes. The interaction was not significant either (*F*(2,30) = 1.983, *p* = 0.155). The same analysis applied on the Letter cancellation test also revealed a significant main effect of group (*F*(1,13) = 8.624, *p* = 0.012), indicating again that non-neglect patients on average found more targets per time (*M* = 0.278, *SD* = 0.090) than patients with neglect (*M* = 0.140, *SD* = 0.091). The main effect of screen size was significant as well (*F*(2,26) = 5.219, *p* = 0.012); the interaction was not significant (*F*(2,26) = 0.176, *p* = .839). According to post hoc comparisons (Fisher’s Least Significant Difference), more targets per time were found in condition TS large (*M* = 0.235, *SD* = 0.120) than in condition TS small (*M* = 0.183, *SD* = 0.066, *p* < 0.05).

**Figure 5.**
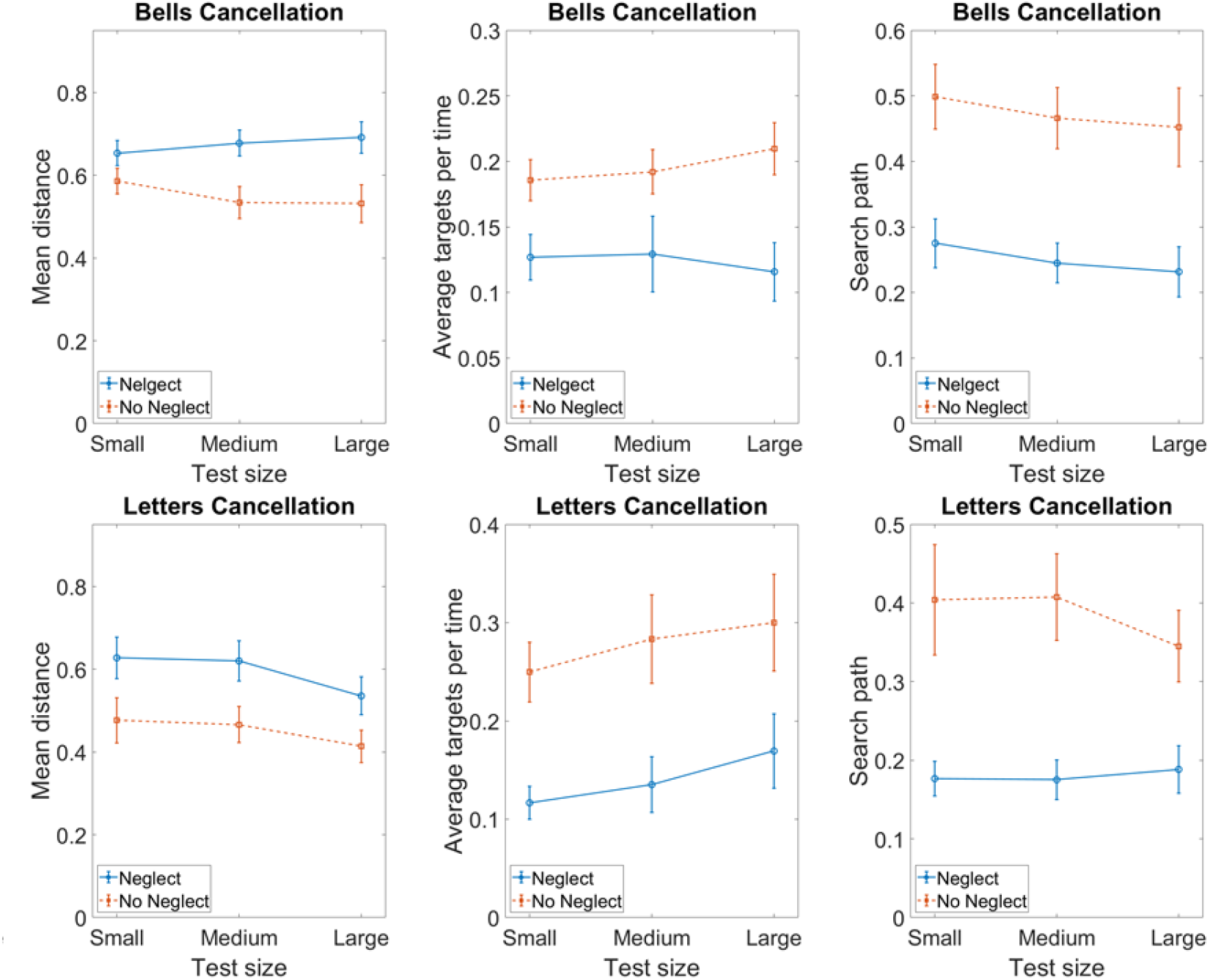
Additional search parameters over different test sizes. Averaged mean distance between to targets found in direct succession (*left panel*), averaged number of targets identified per time (*middle panel*), and averaged mean distance between two successive targets (*right panel*) (including standard error) in the three different touch screen formats of the digitalized cancellation tests in patients with and without spatial neglect (Neglect, No Neglect).

##### Mean distance

Data are illustrated in Figure 5. Analysis of the Bells test revealed that the main effect of screen size (*F*(2,30) = 0.250, *p* = 0.781) and the interaction (*F*(2,30) = 3.109, *p* = 0.059) were found to be not significant, while the main effect of group was significant (*F*(1,15) = 7.357, *p* = 0.016). Apparently distances between two consecutively found targets were smaller for right control patients (*M* = 0.550, *SD* = 0.093) than for patients suffering from neglect (*M* = 0.674, *SD* = 0.093). For the Letter cancellation test, significant results were obtained for both the main effect of group (*F*(1,13) = 5.486, *p* = 0.036) and of screen size (*F*(2,26) = 5.528, *p* = 0.010). The interaction was not significant (*F*(2,26) = 0.244, *p* = 0.785). Again, distances between two consecutively found targets were smaller for right control patients (*M* = 0.452, *SD* = 0.116) than for patients suffering from neglect (*M* = 0.594, *SD* = 0.115). Post-hoc comparisons indicated that in the TS large version (*M* = 0.474, *SD* = 0.120) items found in direct succession were on average closer to one another than in the TS small (*M* = 0.552 *SD* = 0.143*, p* = 0.026) and TS medium (*M* = 0.543, *SD* = 0.127, *p* = 0.018) versions.

##### Search strategy

Data are illustrated in Figure 5. Analysis of this parameter for the Bells test did neither show a main effect of screen size (*F*(2,30) = 2.503, *p* = 0.099) nor an interaction (F(2,30) = 0.002, *p* = 0.998), while the main effect of group was significant *F*(1,22) = 9.11, *p* < 0.01). Control patients without neglect scored significantly higher (*M* = 0.472, *SD* = 0.119) than neglect patients (*M* = 0.251, *SD* = 0.117), indicating that search behavior of neglect patients was more strategic than the one of the right hemispheric controls. Results concerning the Letter cancellation task uncovered a significant main effect for group (*F*(1,13) = 15.787, *p* = .002), indicating that neglect patients search behavior was significantly more strategic (*M* = 0.180, *SD* = 0.51) than the one of right hemispheric control patients (*M* = 0.386, *SD* = 0.101). No significant results were obtained for the main effect of screen size (*F*(2,26) = 0.513, *p* = 0.604) and the interaction (*F*(2,26) = 0.1.177, *p* = 0.324).

### 3.3 Prediction of spatial neglect by the additional parameters of search behavior

While for measuring the CoC variable the test has to be completed, the variables targets per time, mean distance, and search strategy might become apparent early on and might be extractable, even if a test is aborted. For this, however, it is first of all necessary to check whether those new parameters prove to detect neglect specific behavior. To analyze if the three additional parameters of search behavior provided by the digital format can be used to differentiate between right-hemispheric patients with and without spatial neglect, Support Vector Machines (SVM) were used. First, the binary diagnosis (neglect vs. no neglect) was predicted separately for the Bells and the Letter cancellation tests across all screen sizes, using Leave-One-Subject-Out Cross-Validation. Results showed that this cross-participant classification across screen sizes was highly accurate for the Bells test and for the Letter cancellation test with average balanced accuracy scores of 97.92% and 88.19%, respectively. The training and test accuracies for all models are shown in Figure 6. Second, chi-square tests of independence indicated that the frequency of accurately predicted neglect and right-hemispheric control patients (see Table 2) was independent of the screen size for the Bells (χ^2^(2) = 0.05, *p* = 0.973) and Letters cancellation task (χ^2^(2) = 0.18, *p* = 0.914).

**Table 2.** Frequency of correctly classified neglect and righ-hemispheric control patients by screen size (TS small, TS medium, and TS large)

| Screen size | Diagnosis | Prediction Accuracy |  |
| --- | --- | --- | --- |
|  |  | Bells cancellation test | Letters cancellation test |
| TS small | control (n = 8) | 7 | 7 |
|  | neglect (n = 12) | 12 | 9 |
| TS medium | control (n = 8) | 8 | 7 |
|  | neglect (n = 12) | 12 | 12 |
| TS large | control (n = 8) | 8 | 7 |
|  | neglect (n = 12) | 12 | 9 |

**Figure 6:**
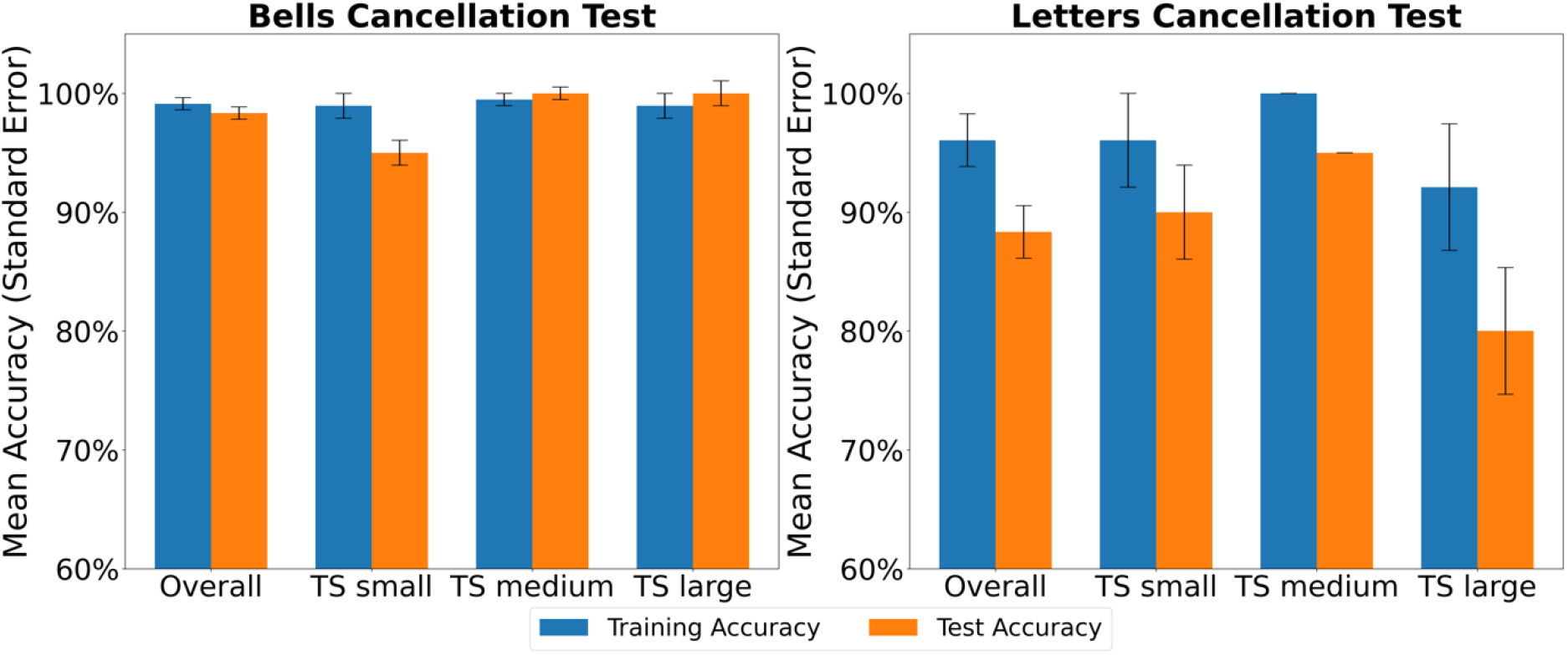
Machine learning model performance. Training and test performance of the machine learning models classifying the binary diagnosis “neglect vs. no neglect” in the right hemispheric patient sample overall and broken down by screen size (TS small, TS medium, and TS large).

## 4. Discussion

### 4.1 Paper-and-pencil vs. digital test version

Several papers have acknowledged numerous perks of digitizing neuropsychological assessments in general (Bauer et al., 2012; Germine, Reinecke, & Chaytor, 2019) and neglect diagnostics specifically (Donelly et al., 1999; Bonato, Priftis, Marenzi, Umiltà, & Zorzi, 2012; Bonato & Deouell, 2013). This has inspired the introduction of novel computer-based neglect assessments (Donelly et al., 1999; Bar-Haim, Kizony, Shahar, & Katz, 2006; Bonato, Priftis, Umiltà, & Zorzi, 2013; Dalmaijer et al., 2015; Villareal et al., 2020). Digital versions have been argued to be more felxible, allowing e.g. to create several parallel versions of a specific task and therefore to prevent learning effects from numerous repetitions of one identical version, e.g. in the course of rehabilitation (Bonato & Deouell, 2013). Moreover, it has been put forward that digital formats could further increase a test’s sensitivity by increasing the amount of information extractable from the test’s data (Bonato & Deouell, 2013, Dalmaijer et al., 2015). However, previous studies also have stressed the importance of test validation digital formats (Bauer et al., 2012; Germine et al., 2019). The present paper is to our knowledge the first that systematically compared patients’ performances between digital and analogous formats. Patients’ CoC scores did not vary considerably between paper-and-pencil versus digital versions. Thus, it seems safe to introduce digitized diagnostic measures (at least in the scope of size variations as investigated in the present study) and keep the existing cut-off scores, without having to fear distortions in the CoC and related diagnostic decisions. The CoC derived from cancellation tasks seems robust to test-digitization.

### 4.2 Test/display size of the cancellation tasks

#### 4.2.1 Center of Cancellation

Patients’ CoCs also did not seem to be impacted by test size. Our analysis revealed that neglect patients ignored a comparable ratio of contralesional target stimuli, regardless of test size. This observation corresponds to previous findings on reference frames suggesting a dynamic view of the neglected area in space, depending on the respective behavioral goal of the subject. Karnath and Niemeier (2002) argued that the brain continuously organizes and re-organizes the representation of the same physical input according to the changing task requirements. The authors showed that whether or not neglect patients ignored certain space in a visual search task did not depend on the frame size itself, but rather on the relative location within the part of space they were asked to pay attention to. Patients were found to ignore the left half of space when asked to explore only that very segment, but attended to it fully when it constituted the right half of a larger segment. Similar results were found by Baylis and colleagues (2004). Further support for this notion has been the observation that removing targets once they are identified by patients reduces patient’s attention frame and thus manages to draw patients’ attention further into contralesional space (Mark, Kooistra, & Heilman, 1988; Keller, Volkening & Garbacenkaite, 2015). In conclusion, these findings indicate that the neglect specific egocentric bias seems to be robust to variations in screen size and provides a suitable explanation for the CoC’s indifference to size changes observed in the present study.

#### 4.2.2 Additional parameters of search behavior

With respect to the additional parameters that became available through digitization, neglect patients showed slower search behavior and increased distance between two consecutively identified items than right hemispheric control patients without neglect in both cancellation tasks. These disadvantages do not seem surprising. It is a frequent clinical observation that neglect patients tend to start working on tasks from the right side and effortfully drag their attention towards the contralesional hemispace. An eye-tracking study on reading behavior, for example, illustrated how straining it is for a neglect patient to advance further towards the neglected left. While a healthy reader finds the beginning of the next line of a text by performing long, pointed saccades, the investigated neglect patient moved leftward gradually (Karnath & Huber, 1992). Of course, such behavior is much more time-consuming. Studies investigating the visual scanning and exploration behavior of neglect patients on photographs (Ptak, Golay, Müri, & Schnider, 2006; Machner et al., 2012; Kaufmann et al., 2020) and videos (Machner et al., 2012) found that increased salience due to motion (Ptak et al., 2006) or contrasts (Machner et al., 2012) can help a patient attend to the neglected hemispace. Kaufmann and colleagues (2020) found neglect patients’ perseverance to ipsilesional space under neutral conditions to be so distinct that it proved to be more sensitive in detecting neglect than common diagnostical measures. This exploration pattern might directly translate to our visual search tasks. Neglect patients may be more likely to move leftward inefficiently progressing from one stimulus to the next, coming across a target every now and then rather coincedentally. This process also makes them more prone to miss a target if a distractor happens to be closer to the current fixation and attracts the patient’s attention instead, resulting in a larger distance between one identified target and the next.

Screen size-dependent performance was found in the Letter cancellation task but not the Bells test, which could be caused by the different complexity of the tests. Neglect according to the Letter cancellation task is diagnosed if more than four contralesional stimuli are omitted, while a diagnosis based on the Bells test requires at least five omissions (Rorden & Karnath, 2010). Out of context, that doesn’t seem a grave difference, however, the Letter cancellation test contains 60, Bells test only 35 targets hidden among distractors. Relatively speaking, a cut-off of 6 % omissions is contrasted to a cut-off of approximately 14 %, indicating that healthy individuals appear to be more likely to omit bells than “A”s. A reason for this might be that automatized letter recognition is known to be superior to object identification (Denckla & Rudel, 1974). This might also explain why in the Letter cancellation but not the Bells test targets were identified faster in the large screen size than the small one. Since automatized reading depends on how well letters are recognizable, the enlarged letters in our paradigm likely improved participants’ perception and search efficiency by making “A”s better distinguishable from distractors. The more effortful shape identification of bells might not benefit as much from a larger depiction. Patients showed significantly faster search behavior only between the small and the large variant of the letters test. Their search was more thorough in the large versionthan in the small and the medium ones when normalized for screen size. These results are hardly surprising, since the size difference between medium and large (59 %) is greater than that between small and medium (28 %). An increase by 59 % reduces the likelihood of missing targets in close proximity to the current target, thus reducing mean distance, while an increase in search speed seems to require a larger increase in test size.

In contrast to the fairly straight-forward measures for search speed and distance to the next target, it is rather hard to come up with a universal indicator of search strategy, since a strategic search can be perfomed in many different ways. The measure we definded as ‘search strategy’ in the present study should cover strategies typically applied by healthy individuals (cf. Fig. 2; Warren, Moore, & Vogtle, 2008). Interestingly, we found that neglect patients’ search behavior was more ‘strategic’ (according to our definition) than that of right hemispheric control patients in both Bells test and Letter cancellation task. While the neglect patients more frequently applied strategies such as those shown in Figure 2, the right hemispheric controls either searched in a less ‘strategic’ manner (according to our definition) or applied a strategy different to the ones typically applied by healthy individuals (Warren et al., 2008). While this finding might seem surprising at first glance, previous research provided evidence that neglect patients do not generally exhibit impairments in search strategy (Donnelly et al., 1999; Mark et al. 2014; Ten Brink et al., 2018). Donnelly and colleagues (1999) generated 16 different strategic search patterns and investigated which ones were applied by healthy participants as well as by right hemispheric patients with and without neglect. Although neglect patients tended to apply different search strategies than the majority of healthy participants and non-neglect patients, their pattern matched some of the authors’ pre-defined strategic search paths. More specifically, neglect patients most frequently applied a strategy that mirrored the most common strategy used by healthy controls. Since our measure accounted for strategies starting from both sides (left or right), both directions were considered as equally strategic.

With regards to the diagnostic use of the three process measures (i.e., time per target, mean distance between targets, and search strategy), machine learning classifications indicated that these variables can be used to differentiate neglect patients from right hemispheric controls reliably, using parsimonious modelling approaches. Particularly for the Letter cancellation test, the overall small differences between training and test accuracy for these cross-participant classifications (see Fig. 6) indicate that the models generalize well, which is crucial for potential applications of such measures. For instance, digital tests could capture these measures in real time and predict the diagnosis already at early stages of the testing procedure, which would be beneficial if, for example, the test has to be aborted. This, in turn, could serve as basis for adaptive and time-saving diagnostic procedures that are less strenuous for patients. While overall predictions were highly accurate in both tests (97.92% for the letter cancellation test and 88.19% for the bells cancellation test), it is important to note that predictions for the letters cancellation test predictions were less accurate and showed larger variation between training and test samples than for the bells cancellation task. Here, future research with larger, balanced samples should investigate whether there are significant differences in the diagnostic value of processes measures between tests. With regards to screen size, our analyses showed that model predictions for both tests were independent of screen size. For further studies and potential practical applications, this indicates that additional measures obtained from digitized tests can be used to reliably classify neglect regardless of screen size. Nonetheless, future research with larger sample sizes is required to confirm the robustness of our models.

## 5. Conclusion

The present results allow an optimistic outlook on the digitization of cancellation tasks. A change in modality and size does not seem to bias a patient’s CoC, which often serves as the basis for diagnostic decision-making. This robustness to the diagnostic outcome opens the possibility to optimize some visual parameters for more efficient testing. Increasing the stimulus size in the Letters cancellation test seems to help patients identify targets more quickly, which would make diagnosis less time consuming for the examiner and less exhausting for the patients. Machine learning methods indicated that new search parameters derived from computerized tests could help differentiate neglect and non-neglect patients. The latter is an interesting new perspective, because in clinical practice it is often not possible to perform the cancellation tasks to the end (which is mandatory to calculate the CoC measure). If performance features charaterisic for spatial neglect can be extracted from variables such as targets per time, mean distance, and search strategy and might become extractable already early on during test execution, neglect diagnosis might become possible even if the test is discontinued. Future studies are needed to investigate the latter.

## Acknowledgements

Funding: This work was supported by the Deutsche Forschungsgemeinschaft (KA 1258/23-1).

## Notes

http://dx.doi.org/10.17632/6dzxs69j7d.1

### Competing Interest Statement

The authors have declared no competing interest.

